# Landscape heterogeneity, forest structure and mammalian host diversity shape *Ixodes ricinus* density and *Borrelia* prevalence in German forests

**DOI:** 10.1101/2025.05.13.653726

**Authors:** Sara Weilage, Max Müller, Lara Maria Inge Heyse, Dana Rüster, Manfred Ayasse, Martin Pfeffer, Anna Obiegala

## Abstract

Ticks, particularly *Ixodes ricinus*, and the associated Lyme borreliosis risk, represent key concerns within the One Health framework, prompting extensive research in this field. However, comprehensive studies that jointly consider landscape characteristics, local forest structure and management, climate, and host community composition – alongside direct measures of tick density and *Borrelia* spp. infection status - are scarce.

In this study, we test the hypothesis that habitat diversity exerts a dilution effect, primarily by supporting greater diversity of mammal hosts. Therefore, we examined *I. ricinus* tick density and *Borrelia* spp. prevalence in relation to a comprehensive set of habitat and host-related variables. Ticks were collected using the flagging method and mammal hosts were monitored using an innovative camera-trapping approach across 25 forest plots along a land-use gradient within the Schwäbische Alb exploratory in Germany. Our findings indicate that both tick density and *Borrelia* spp. prevalence are influenced by a complex interplay of habitat factors across different spatial scales, as well as the mammal host community composition. Overall, our results provide novel support to the dilution effect hypothesis, suggesting that greater habitat and host diversity contribute to a reduced Lyme borreliosis risk in this region.

## Introduction

Ecological factors play a pivotal role in driving pandemics and disease outbreaks, while also influencing the transmission dynamics of various zoonotic infectious diseases, such as Lyme borreliosis. In North America, a relatively well-defined host-vector-pathogen relationship, primarily involving the white-footed mouse (host), *Ixodes scapularis* (vector), and *Borrelia burgdorferi* sensu stricto (pathogen), are the components driving the Lyme borreliosis risk. In contrast, the European ecological transmission cycle of Lyme borreliosis is markedly more complex, involving a broad range of host species (mainly small mammals) across different trophic levels and ecosystems (Mannelli *et al*., 2012, Estrada-Peña *et al*., 2024). Further, Lyme borreliosis in Europe is caused by bacteria from the *Borrelia burgdorferi* s.l. complex, consisting of at least 11 genospecies targeting multiple organic systems with varying severities of clinical outcomes in humans (Rizzoli *et al*., 2011). While host species are found in various ecosystems, the primary vector in Europe, *I. ricinus*, is known to have a high ecological plasticity, thriving in all kinds of habitats through parks, forests, and also different altitudes (Kahl & Gray, 2023). Further, *I. ricinus* is a major vector in Europe, capable of transmitting a wide range of zoonotic pathogens. This capacity is largely due to its broad host range, encompassing over 300 vertebrate species, which are essential for the tick’s developmental life cycle (Gern & Humair, 2002). Forest management, landscape configuration, climate, and host community composition all affect tick density and *Borrelia* spp. infection rates in ticks (Ehrmann *et al*., 2018, Bourdin *et al*., 2022, Król *et al*., 2022, Boulanger *et al*., 2024). These factors are interdependent, yet they are rarely assessed collectively and can be challenging to quantify especially in the complex European host-vector-pathogen system. Consequently, while the dilution effect hypothesis has gained empirical support in North America, suggesting that biodiversity loss can increase disease risk by elevating the relative abundance of competent reservoir hosts (Allan *et al*., 2003), its applicability in European systems remains uncertain. The greater host and pathogen diversity, coupled with varied forest management practices, complicates generalization. Indeed, European studies have yielded mixed (Gandy *et al*., 2022) or even contradictory (Ruyts *et al*., 2018) results, highlighting the need for more comprehensive approaches.

The Swabian Alb in southwestern Germany exemplifies a region of high biodiversity and forest management gradients and thus offers an ideal setting to investigate these interactions. A previous study in the Swabian Alb suggested that on the one hand forest management practices have an important impact on questing *I. ricinus* nymph density but on the other hand that variable weather conditions may override these effects (Lauterbach *et al*., 2013). However, in the latter study only a small size of individuals was sampled, the diversity of mammal hosts was not considered and the pathogenic potential of the ticks has not yet been assessed. Numerous other European studies have also examined the effects of specific variables on *I. ricinus* density or *Borrelia* prevalence (Ehrmann *et al*., 2018, Estrada-Peña *et al*., 2024, Olsthoorn *et al*., 2025). However, only a few have investigated the combined influence of habitat and host-related factors (Halos *et al*., 2010, Ruyts *et al*., 2018, Gandy *et al*., 2022, Boulanger *et al*., 2024), and these studies still lack a fully comprehensive approach.

Drawing on this existing literature, we aimed to test the hypothesis that habitat diversity across different scales exerts a dilution effect, primarily by supporting greater diversity of mammal hosts, with small mammals playing a key role. To achieve this, we made use of the comprehensive set of habitat variables provided by the Biodiversity Exploratories database and supplemented it with targeted data collection of mammal hosts. Thus, to our knowledge, this is the first study to integrate landscape characteristics, forest structure and management intensity, climate, and mammal community host composition to assess their combined effects on *I. ricinus* density and *Borrelia burgdorferi* s.l. prevalence in central Europe.

## Material & Methods

### Study Region

The Swabian Alb, located in the Reutlingen district of Baden-Wurttemberg, is one of three designated long-term research areas in Germany for biodiversity and ecosystem studies within the Biodiversity Exploratories programme (Fischer *et al*., 2010). With an area of 420 km² it spans altitudes ranging from submontane to montane zones. The landscape is a patchwork of forests and open land, reflecting a variety of land-use practices. Since 2007, the Swabian Alb has been the focus of forest and grassland experiments, as well as extensive ecological observations. Data collected from the region, covering both biotic and abiotic factors, are accessible through an online platform and are integral to long-term scientific analysis (Chamanara *et al*., 2021).

For this research, 25 of the 50 forest plots available in the Swabian Alb were chosen, representing the full range of the land-use gradient which was assessed using the Silvicultural Management Intensity Index (SMI), which evaluates key characteristics such as stand age, stand growth, and dominant tree species (Schall & Ammer, 2013) (Figure 1). In our case, lower SMI values indicate more extensively managed, beech-dominated forests, whereas higher SMI values correspond to more intensively managed stands dominated by conifers.

**Figure 1:**
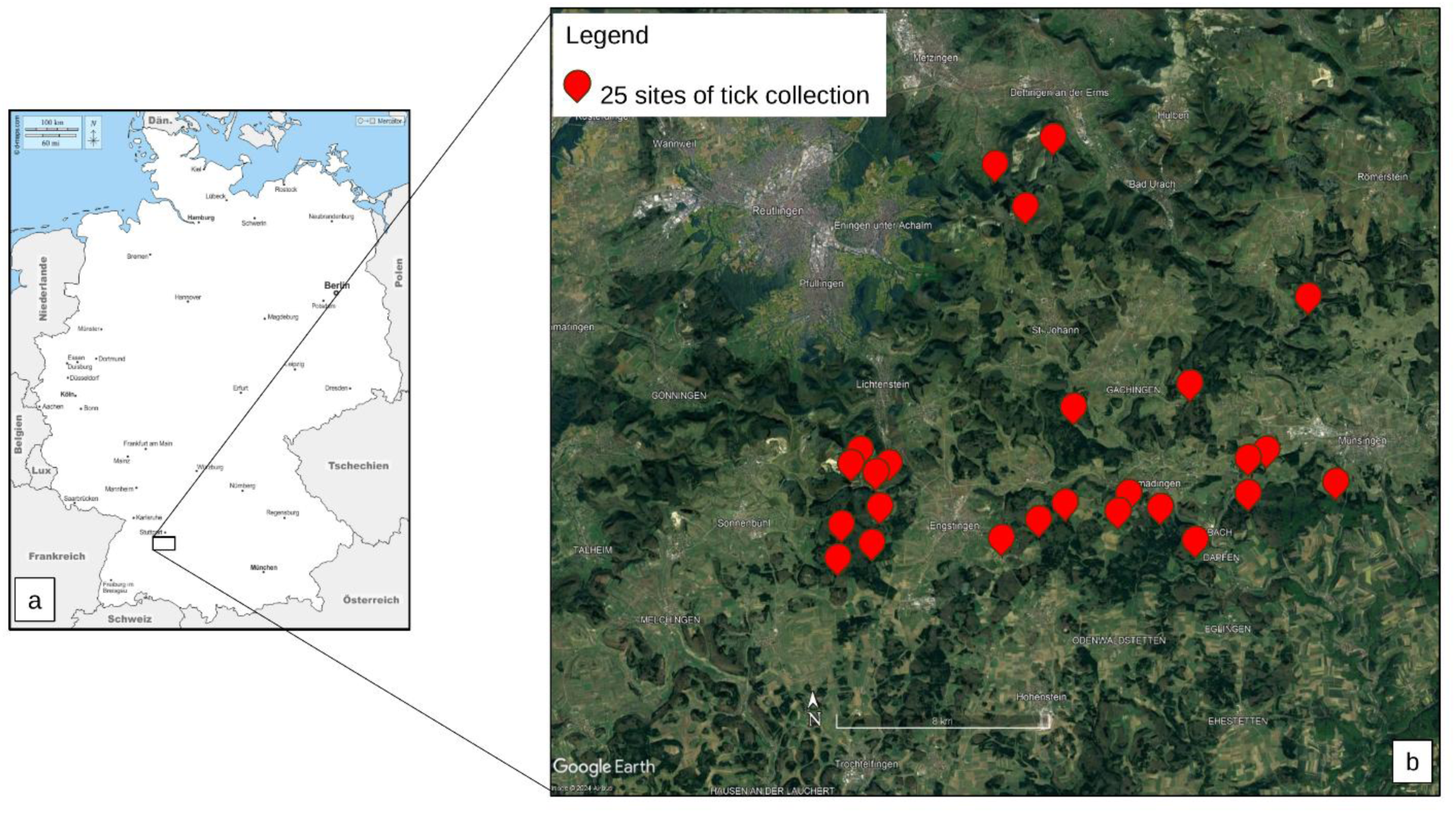
Geographical location of 25 tick collection sites (marked in red) along a land-use gradient in the Swabian Alb, Germany. **a** shows the location of the study area in Germany and **b** the location of every single plot. The image was created using Google Earth Pro, Map: Google Earth ©2025 Google, Image © 2025 Airbus, Image © 2025 Maxar Technologies and https://d-maps.com/carte.php?num_car=2002&lang=de.

### Large mammal camera trapping

Two camera traps (Dörr Snapshot Mini Black 30 MP 4K, sporadically predecessor models) were installed on each of the 25 plots according to a common scheme: height ∼50 cm, inclination of 5° downwards, direction north, 3 pictures per trigger, 30 sec non-reactive between triggers (Kammerle *et al*., 2018, Bubnicki *et al*., 2019). Two cameras per plot minimize site effects and prevent data gaps in the event of camera failure. Traps were installed at suitable locations enabling sufficient range and undisturbed triggering (Wening *et al*., 2019). Cameras were technically maintained at least every 6 months and relocated randomly to further minimize site effects. They were set up during the entire study from spring 2023 to spring 2024 with a total of 9,609 camera days included in the analysis.

Processing of raw images and species identification was done using ‘Agouti’, an AI powered platform for managing wildlife camera trapping projects (Casaer *et al*., 2019). AI-based species identifications (AI tool ‘Western Europe v4a’) were subsequently confirmed through direct, in-person verification. Recordings of human activity were not taken into consideration.

### Small mammal camera trapping

Custom-built small mammal camera traps, adapted from the design by Littlewood *et al*. (2021), were used for small mammal monitoring (Figure 2). For this purpose, Dörr camera traps were equipped with a 4x close up-lens (dHD Digital), the flash unit was adjusted to a lower power level to prevent overexposure and attached to a box baited with a mixture of peanut butter, oats and apples. The bait was placed out of reach under a grid to exclude a reward for the animal and to minimize frequent returns of the same individual. Small mammal camera trappings took place at the same location and during the same time periods as tick collection, as described below, in summer and autumn 2023 and in spring 2024. Two traps per each 300 m^2^ area were set up for seven days per season which equals a total of 984 camera days. To facilitate species identification, traps were set to record 15 s videos at each trigger with 30 s delay before they could be triggered again. Species were identified according to nature field guides (Braun & Dieterlen, 2003). In the case of *Apodemus*, species identification of *A. flavicollis* or *A. sylvaticus* was possible in 86% of the recordings. The rest were only identified to genus level and subsequently not used for Shannon diversity calculations.

**Figure 2:**
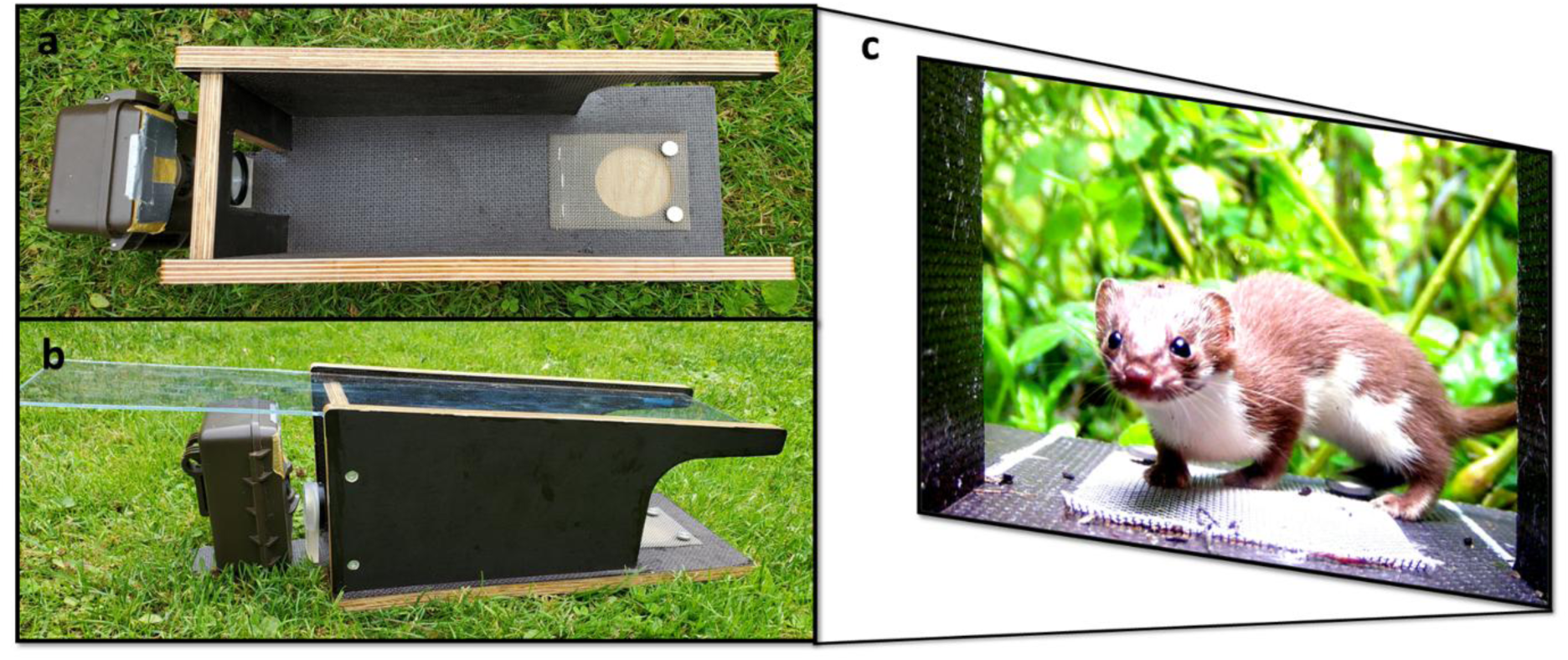
Small mammal camera trap adapted from Littlewood *et al*. (2021). Top and side view of Dörr Snapshot Mini Black 30 MP 4K extended by a 4X magnifying lens, dimmed LED flash and box with bait under grid (**a**, **b**). Still image of a video (**c**, *Mustela nivalis*).

### Tick collection, species identification and density calculation

In spring (May), summer (August), and autumn (October) of 2023, as well as in spring (May) of 2024, ticks were collected using the flagging method with a 100×100 cm white wool-cotton cloth and stored separately for each location at −20°C until further examination. The sampling area at each of the 25 previously mentioned plots measured 12×25 m and was permanently marked resulting in a total flagged area of 300 square meters per plot. Ticks were assigned to their developmental stage and taxonomically identified to species level by using morphological keys (Estrada-Peña *et al*., 2017) under a stereomicroscope (Motic® SMZ–171, Motic Europe, S.L.U., Barcelona, Spain). The nymphal tick density per 100 m^2^ was calculated for each location. Subsequent analyses were conducted exclusively on nymphal ticks.

### DNA extraction of ticks and molecular analyses for *Borrelia* spp

To prepare DNA purification, 1 g of 2.8 mm steel beads (Peqlab Biotechnology, Erlangen, Germany) and 200 µl of PBS were added to each individual tick sample. Subsequently, each sample was homogenized using a Precellys®24 tissue homogenizer (Bertin Technologies, Montigny Le Bretonneux, France) at 5000 rpm for two cycles of 30 seconds each, with a 15-second pause between cycles. DNA extraction was performed using the QIAamp DNA Mini Kit (Qiagen Germany, Hilden) according to manufacturer’s instructions. The quality and quantity of DNA in each sample were assessed using a NanoDrop spectrophotometer (NanoDrop® 2000c, Thermo Fisher Scientific, Waltham, MA, USA).

The tick samples were screened for the presence of *Borrelia* spp. DNA by detecting the p41-flagellin gene (96 base pairs) through real-time polymerase chain reaction (PCR) using the Mx3000 Real-Time Cycler (Agilent, Santa Clara, CA, USA) in accordance with an established protocol (Schwaiger *et al*., 2001).

### Data availability

This work is based on data elaborated by the Biodiversity Exploratories programme (DFG Priority Program 1374). The datasets are publicly available in the Biodiversity Exploratories Information System (http://doi.org/10.17616/R32P9Q). The datasets are listed in the references section.

However, to give data owners and collectors time to perform their analysis the Biodiversity Exploratories’ data and publication policy includes by default an embargo period of three years from the end of data collection/data assembly which applies to the remaining datasets (IDs: 32109, 32111, 32113 (Müller, 2025, Müller, 2025, Weilage, 2025)). These datasets will be made publicly available via the same data repository.

### Statistical analysis

Relative abundance indeces (RAIs) were calculated for each species as events per 100 camera days in R with the script described in Rovero & Zimmermann (2016):

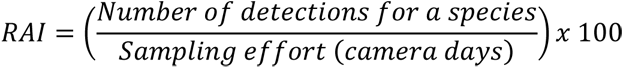

The standardization of detections by the sampling effort makes the results comparable across time, space and methods.

Shannon’s diversity index (H) was calculated from RAIs:

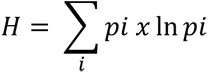

with 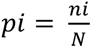, N = total sum of RAIs, n_i_ = sum of RAIs of respective species group. We tested the effects of landscape variables, local forest management and structure, climate variables and the mammal community on nymph density and *Borrelia* spp. prevalence in nymphs (Table 1). For nymph density as response variable we fitted generalized linear models (GLMs). For *Borrelia* spp. prevalence as response variable we fitted linear models (LMs). Due to high collinearity (VIF>10) air temperature at 2 m height, herb cover and conifer share on plot scale and forest edge density on landscape scale had to be excluded from the models. All statistical analyses were conducted in R version 4.3.1 (2023-06-16) (Team, 2023). GLMs and LMs were run using the ‘glm’ and ‘lm’ function from the *stats* package, and the ‘glmTMB’ from the *glmmTMB* package (Brooks *et al*., 2017). We selected and averaged the best fitting models (ΔAICc < 2), using the MuMln package (Barton, 2020). We report conditional averaged model outputs and the full set of candidate models are provided in the Supplementary Information. For the seasonal analysis best models are reported for nymph density as model dredging returned only single top-ranked models (AICc≤2.0). In the seasonal models, total species richness had to be excluded due to high collinearity (VIF>10).

**Table 1:**
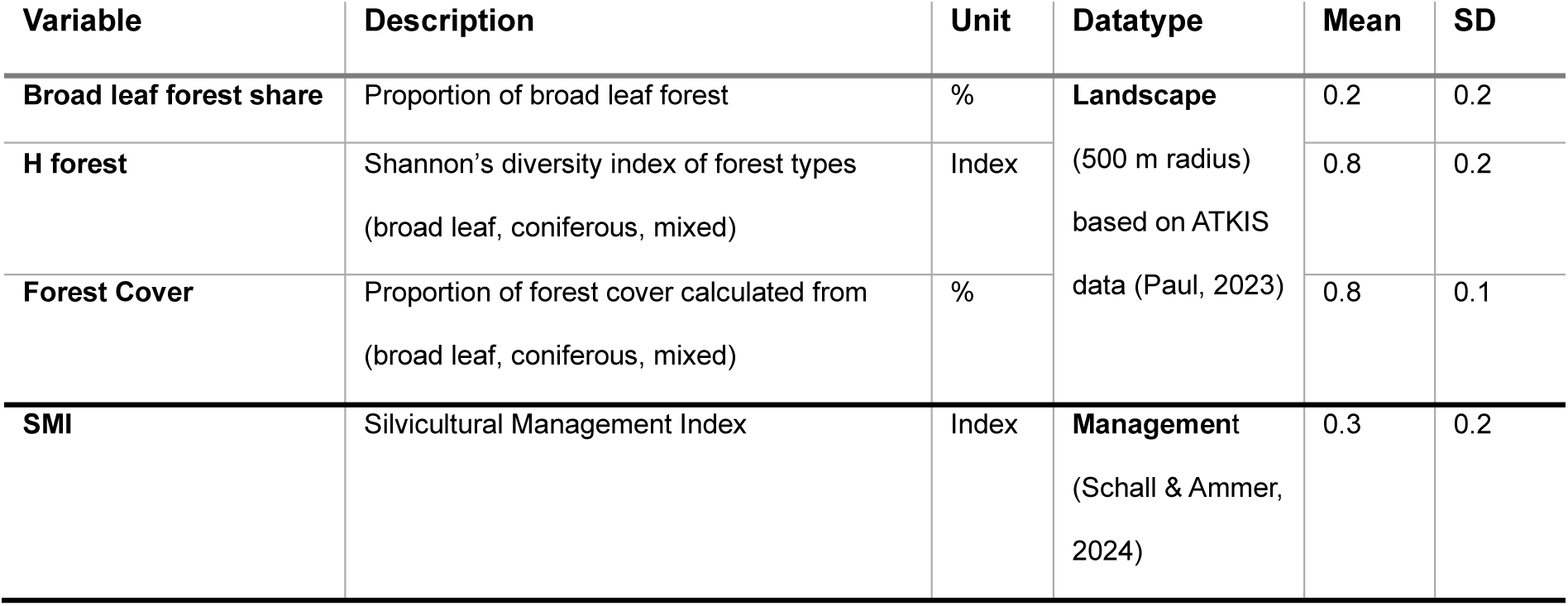
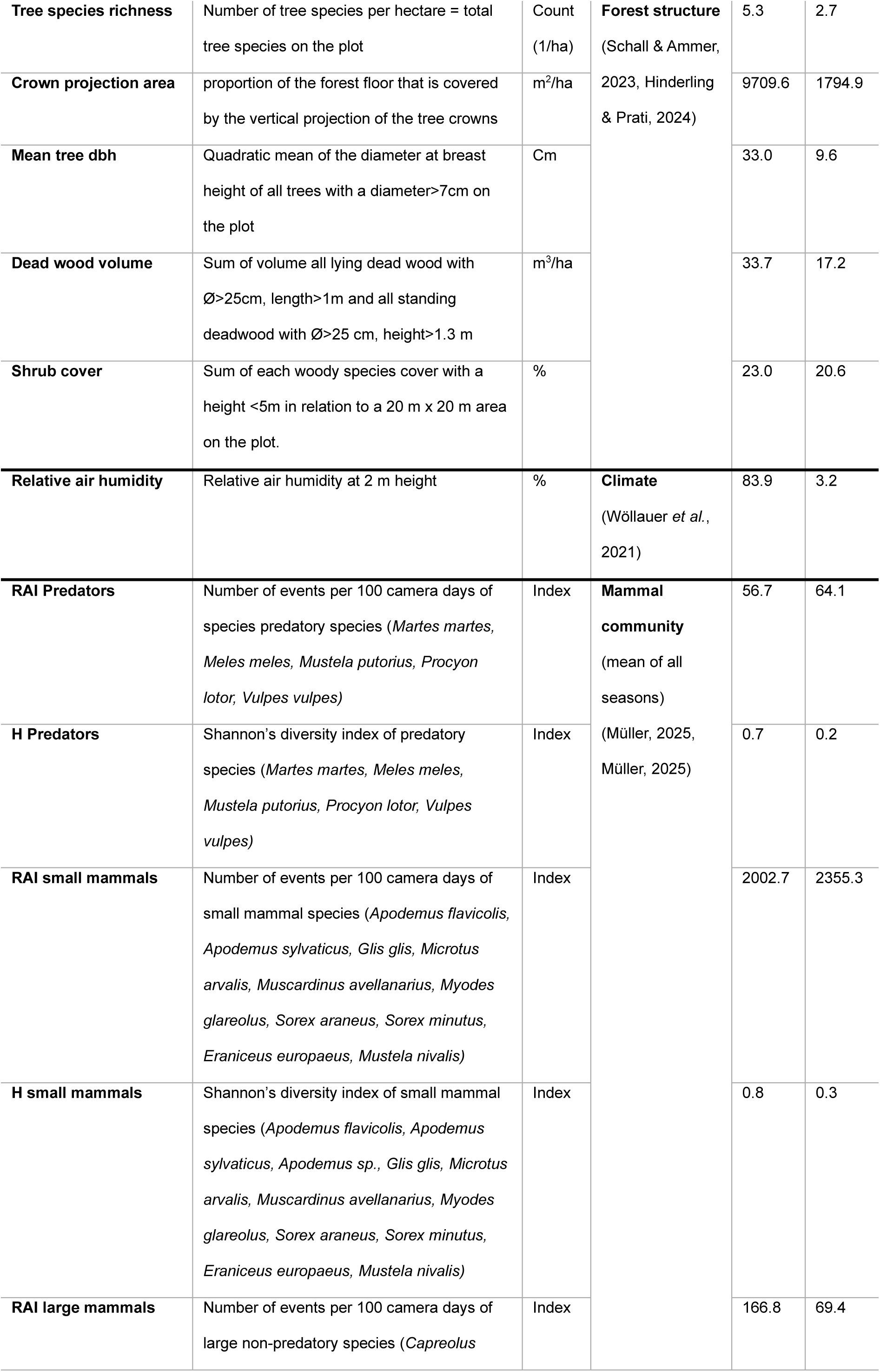
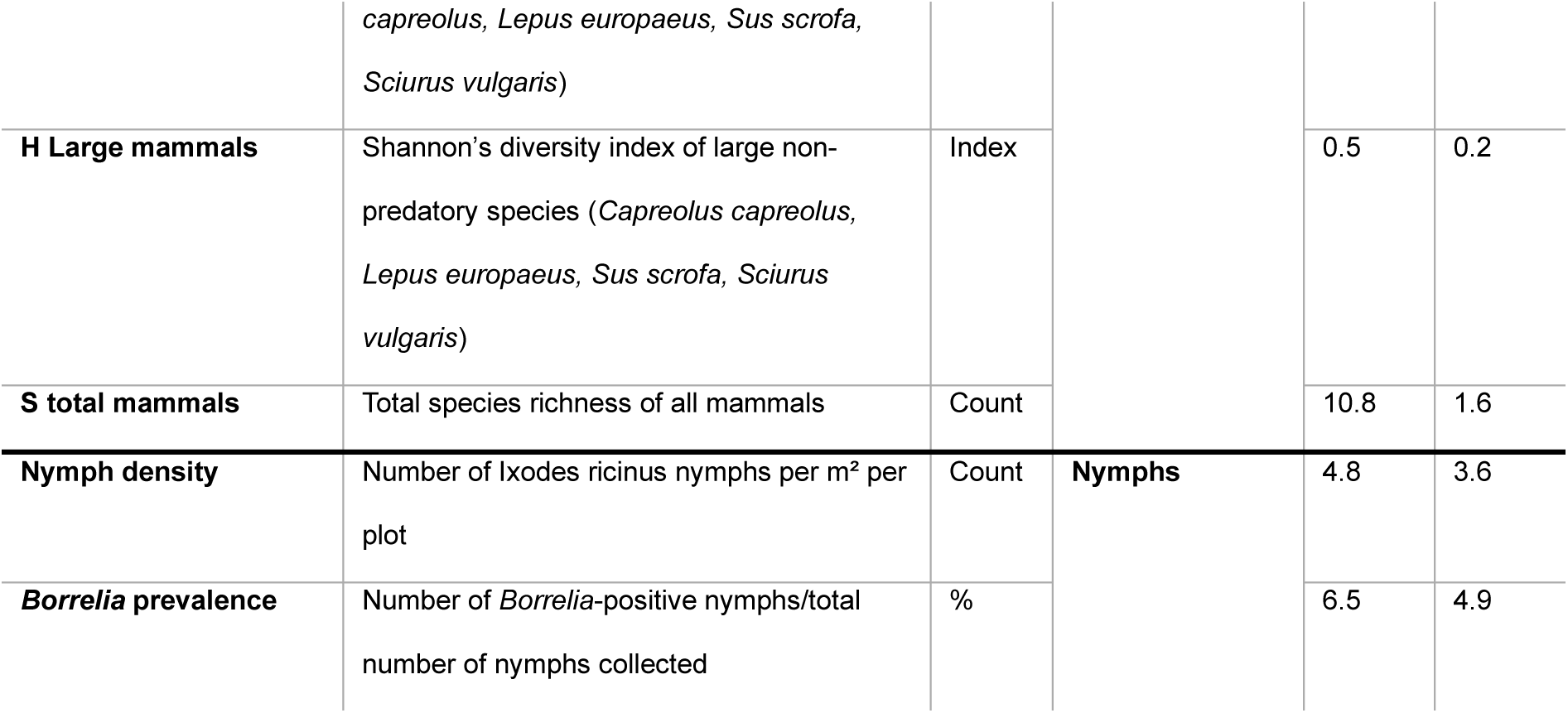
Variables used in the regression models to determine nymph density and *Borrelia* spp. prevalence in nymphs. Landscape variables were calculated within a 500 m buffer around plot centers; local forest structure, management, and mammal diversity metrics were derived from data within the one ha plots.

We hypothesize that different groups of mammals have distinct functions in this predator-prey-host system and therefore have differing effects on the density of *I. ricinus* nymphs and *Borrelia* spp. prevalence. Most larger mammals are considered “dilution” or zooprophylactic hosts (see Table 1, *Capreolus capreolus, Lepus europaeus, Sus scrofa, Sciurus vulgaris*) while most small mammals are considered reservoir hosts for *Borrelia* spp. Larger predatory mammals are considered “maintaining” hosts for *Borrelia* spp. and also regulate small mammal populations, important hosts for *I. ricinus* nymphs (Ruyts *et al*., 2018). However, due to the methodology the red squirrel was recorded with large mammal camera traps and consequently included in this group.

## Results

### Tick collection

A total of 1816 ticks were collected. Two individuals could not be assigned to species level due to their poor conservation status, but all other collected ticks were identified as *I. ricinus* (n = 1814). Nymphs represented the most abundant life stage, with 1439 individuals (79.24%; CI 77.32–81.04), followed by larvae (n = 266; 14.65%; CI 13.09-16.35), females (n = 63; 3.47%; CI 2.72-4.42), and males (n = 48; 2.64%; CI 1.99-3.49) (Table 2, S 1). Most nymphal ticks were collected in spring 2024 (n = 462; 32.11%; CI 29.69-34.52) followed by summer 2023 (n=385; 26.75%; CI 24.47-29.04), spring 2023 (n = 338; 23.49%; CI 21.14-25.64) and autumn 2023 (n = 254; 17.65%; CI) (Table 2).

**Table 2:**
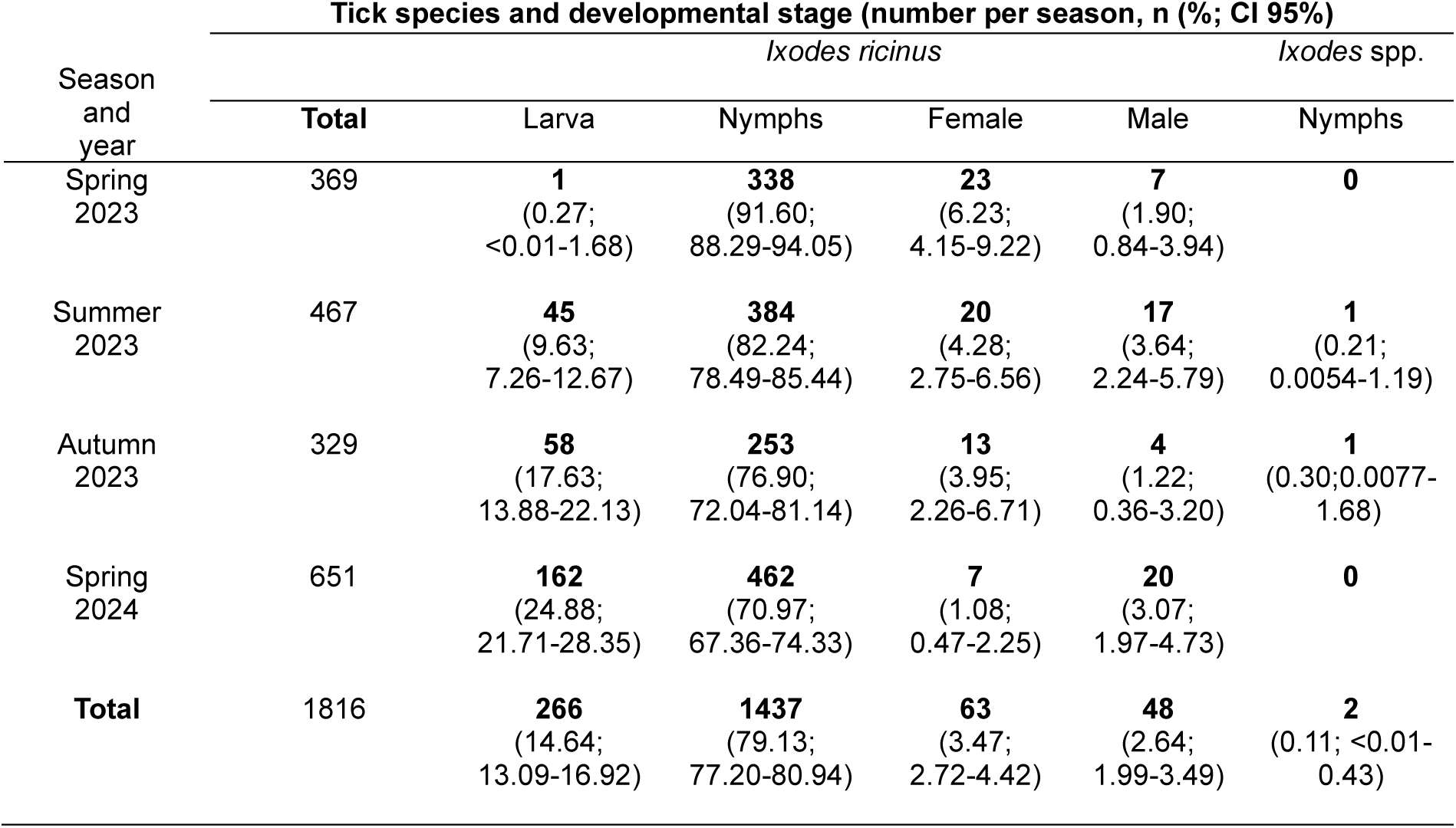
Species, developmental stage and number of collected ticks per season and in total. n = collected ticks (%; Confidence interval (CI 95%))).

Only nymphal developmental stages were included in the analyses of *Borrelia* spp. prevalence, as they occur in higher numbers than adults, exhibit significantly higher *Borrelia* spp. infection rates than larvae, and pose a major risk to humans due to their small size, which makes them difficult to detect (Table 3, S 1).

**Table 3:** *Borrelia burgdorferi* sensu lato prevalence in all tested nymphal *Ixodes ricinus* ticks among the different seasons and in total. n = positive ticks/ tested ticks (%; Confidence interval (CI 95%)).

| Season and year | *Borrelia burgdorferi* sensu lato prevalence |
| --- | --- |
| Spring<br>2023 | **16/338** (4.73; 2.73-7.57) |
| Summer<br>2023 | **22/385** (5.71; 3.63-8.51) |
| Autumn<br>2023 | **24/254** (9.45; 6.61-13.74) |
| Spring<br>2024 | **30/460** (6.52; 4.44-9.24) |
| **Total** | **92/1437** (6.40; 5.19-7.82) |

### Camera trapping

We recorded 5438 large mammal events of four non-predatory and five predatory species over 9609 camera days on the 25 plots from spring to autumn 2023 and in spring 2024 (Müller, 2025). In summer and autumn 2023 as well as in spring 2024, 4626 small mammal events of 10 species were collected in a total of 984 camera days on the same 25 plots (Müller, 2025). These mammal community data were combined with the silvicultural management index (SMI), local forest structure attributes, landscape characteristics and climate variables to test the dilution effect hypothesis (Table 1).

### Ecological determinants of nymph density

Nymphal tick density strongly varied across the 25 forest plots, ranging from 0.00 - 46.67 individuals per 100 m^2^. The conditional average of the generalized linear regression model retained nine explanatory variables, with eight exhibiting statistically significant effects (Figure 3). The density of nymphs was influenced by a combination of landscape characteristics, local forest structure and management practices, as well as the composition of the mammal community (Tables S 2, S 3 and Figure 3).

**Figure 3:**
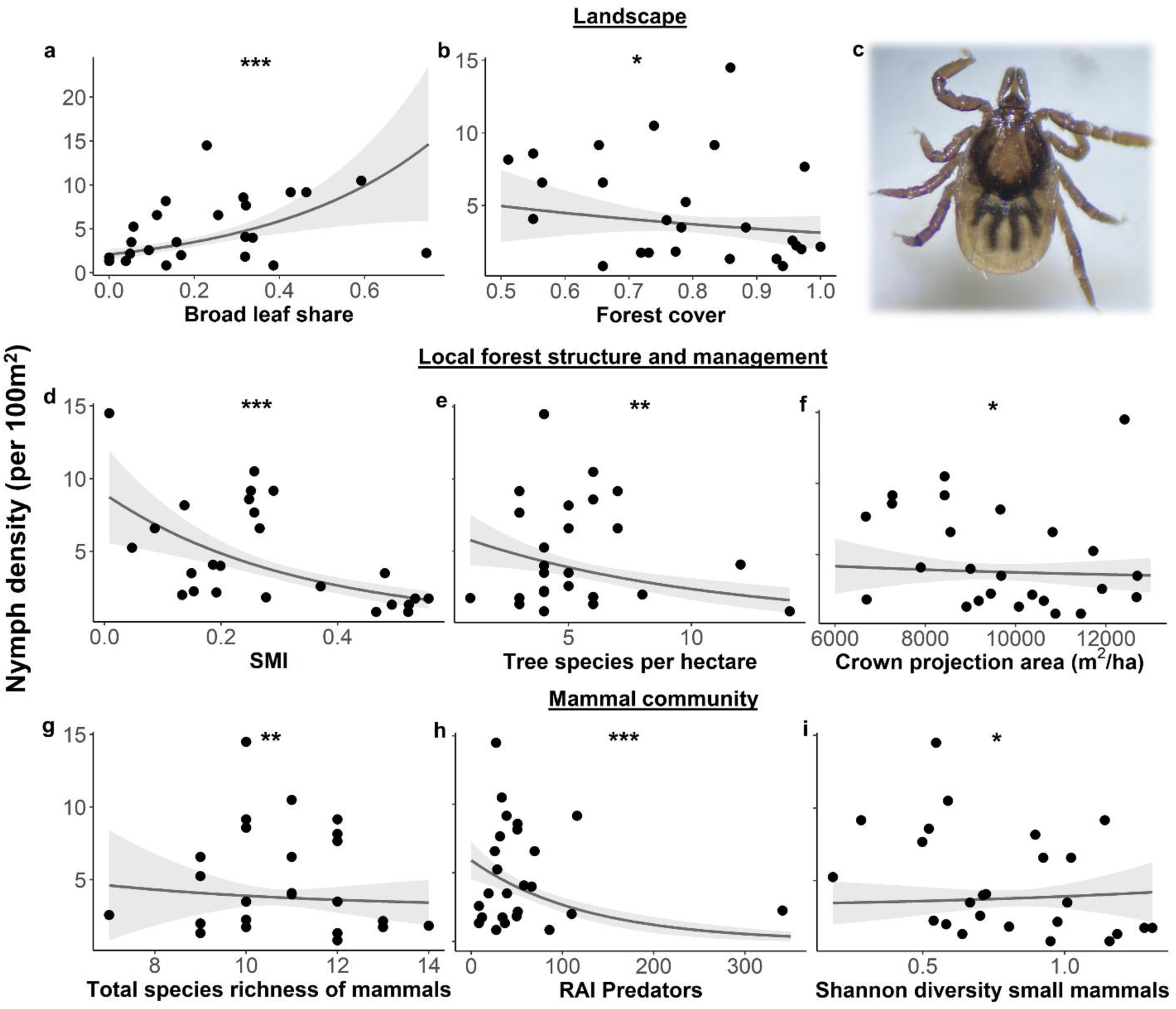
Ecological determinants of nymph density. Nymph density (per 100 m^2^) on 25 forest plots in the Swabian Alb in relation to significant factors of the conditional averaged generalized linear regression model: Landscape factors (broad leaf share, forest cover, **a**, **b**), local forest structure and management (silvicultural management index (SMI), tree species per hectare, crown projection area, **d**, **e**, **f**) and mammal community (total species richness, relative abundance index (RAI) Predators, Shannon’s diversity index small mammals, **g**, **h**, **i**). Scatter points represent data at each forest plot. Lines represent fitted model curves. Shaded areas represent 95% confidence intervals. *p≤0.05, **p≤0.01, ***p≤0.001. Magnified picture of an *I. ricinus* nymph (**c**).

At the landscape scale, nymph density significantly decreased with a lower proportion of broadleaf forest (p<0.001) and with increasing forest cover (p=0.028, Figure 3 a, b). At the local plot scale, higher values of the SMI (p<0.001), greater tree species richness (p=0.003), and — albeit with minimal observable effect within the studied range — a larger crown projection area were all associated with a reduction in nymph density (p=0.017, Figure 3 d, e, f). Regarding mammal community, the RAI of predatory mammals was linked to a decrease in nymph density (p<0.001), while total species richness also contributed to this reduction, albeit with a small observable effect (p=0.001, Figure 3 g, h). Interestingly, the RAI of small mammals showed no evidence of a significant influence on nymph density in this analysis, whereas Shannon’s diversity exerted a small but significant positive effect (p=0.017, Figure 3 i).

### Ecological determinants of *Borrelia* prevalence in nymphs

A total of 1437 nymphal ticks were screened for *Borrelia* spp. Two individuals were excluded from testing as they were used for other analyses. Among the tested ticks, *Borrelia* spp. DNA was detected in 96 specimens; however, applying a cycle threshold (CT) of 41.42 reduced the number of confirmed positive cases to 92, resulting in an overall Borrelia prevalence of 6.4% (CI 5.19-7.82). Prevalence varied substantially across plots, ranging from 0.00 – 19.23%.

The conditional average of the linear regression model retained 10 explanatory variables, eight with statistically significant effects, indicating that *Borrelia* prevalence in nymphs is shaped by a multifactorial interplay of landscape characteristics, local forest structure, climatic factors and the mammal community (S 4, S 5, Figure 4).

**Figure 4:**
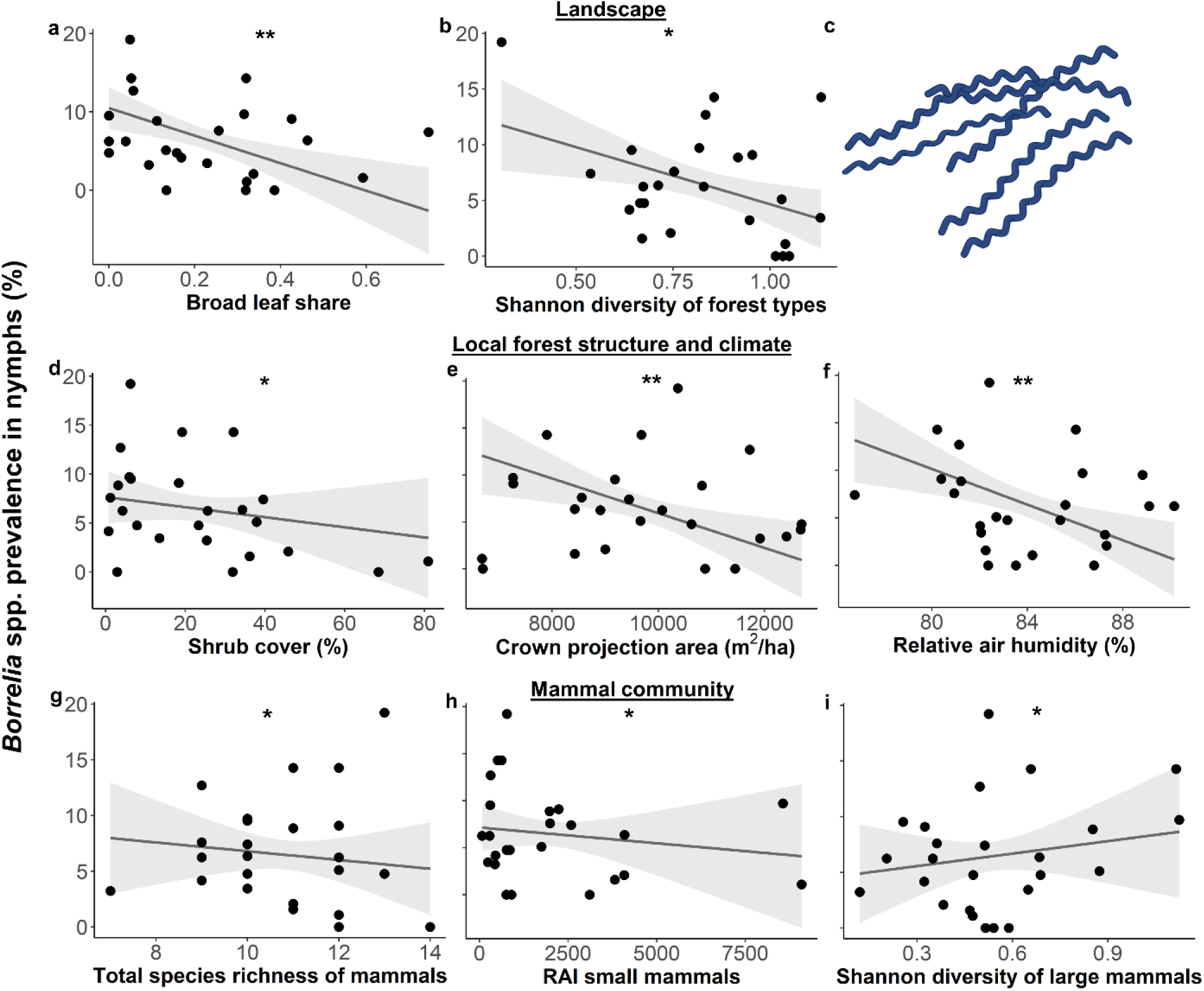
Ecological and climatic determinants of *Borrelia* prevalence in nymphs. *Borrelia* spp. prevalence in nymphs on 25 forest plots in the Swabian Alb in relation to significant factors of the conditional averaged linear regression model: Landscape factors (broad leaf share, Shannon diversity of forest types, **a**,**b**), local forest structure and climate (Shrub cover, Crown projection area, relative air humidity, **d**,**e,f**) and mammal community (total species richness (S), relative abundance index (RAI) of small mammals, Shannon diversity of large non-predatory mammals, **g,h,i**). Scatter points represent data at each forest plot. Lines represent fitted model curves. Shaded areas represent 95% confidence intervals. *p≤0.05, **p≤0.01, ***p≤0.001. **c** Schematic representation of *Borrelia* created with BioRender.com.

At the landscape scale, *Borrelia* spp. prevalence decreased with a higher proportion of broad-leaved forest (p=0.002) and greater diversity of forest types (p=0.012, Figure 4 a, b). On a local scale, higher shrub cover (p=0.032), larger crown projection area (p=0.007), and elevated relative air humidity (p=0.004) were all associated with lower *Borrelia* prevalence (Figure 4 d, e, f). Within the mammal community, both greater total species richness (p=0.017) and higher RAI of small mammals (p=0.014) reduced prevalence (Figure 4 g, h). Notably, however, the Shannon diversity of large non-predatory mammals had a positive effect on *Borrelia* prevalence (p=0.045, Figure 4 i).

### Seasonal effects of the mammal community on nymph density

Since the small mammal population is subject to significant fluctuations, a seasonal analysis of the data was advisable. The development of nymphs after their blood meal as larvae takes approximately three months, making the influence of the mammal population from the preceding season relevant for nymph density and infection risk. To account for this, we analyzed mammal community data collected in one season in relation to nymph density and *Borrelia* spp. prevalence in the subsequent season. Specifically, we examined summer mammal data alongside autumn tick data, representing directly consecutive seasons, and autumn mammal data with spring tick data, spanning the winter period. Nymphal tick densities were ranging from 0.67 - 15.67 per 100 m^2^ in autumn and 0.33 to 46.67 per 100 m^2^ in spring. The best-fitting model retained two explanatory variables for the nymph density in autumn and four variables in spring, respectively. In both seasons, density of nymphs was influenced by a combination of mammal host abundance and diversity (S 6, S 7, Figure 5).

**Figure 5:**
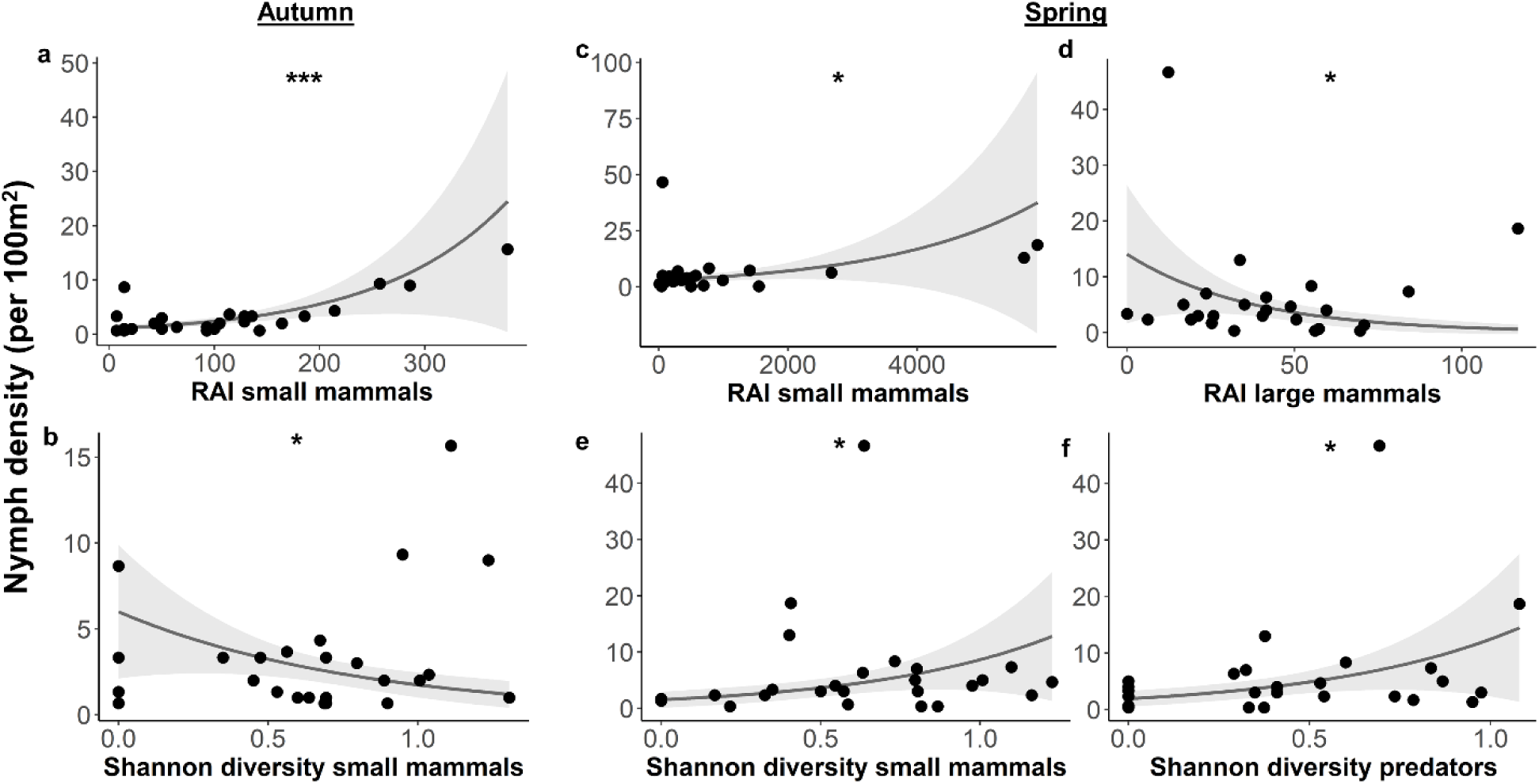
Determinants of the mammal host community for seasonal nymph densities in autumn and spring. Nymph density (per 100 m^2^) on 25 forest plots in the Swabian Alb in relation to significant factors of the best generalized linear regression model in autumn 2023 (**a,b**) and in spring 2024 (**c-f**). Scatter points represent data at each forest plot. Lines represent fitted model curves. Shaded areas represent 95% confidence intervals. *p≤0.05, **p≤0.01, ***p≤0.001.

As hypothesized, nymph density showed a positive relationship with the relative abundance of small mammals in both autumn (p<0.001) and spring (p=0.012, Figure 5 a, c). However, the influence of small mammal diversity differed between the two seasons, showing opposing trends: a negative association in autumn (p=0.014) and a positive association in spring (p=0.020, Figure 5 b, e). Additionally, nymph density in spring was negatively associated with the relative abundance of large mammals (p=0.010) and positively correlated with predator diversity (p=0.011, Figure 5 d, f).

### Seasonal effects of the mammal community on *Borrelia* prevalence in nymphs

A total of 24 nymphs tested positive for *Borrelia* spp. in autumn 2023 and 30 in spring 2024 (Table 3). The highest prevalence was recorded at a single sampling site in autumn 2023 with 50.00% and in spring 2024 with 27.27% (S 1).

The conditional averages retained only one explanatory variable for *Borrelia* spp. prevalence in autumn and three variables in spring. However, the *Borrelia* spp. prevalence in nymphs did not appear to be significantly influenced by the mammal community in either season (S 8, S 9).

## Discussion

Our study takes an integrated approach to understand the ecological drivers of *I. ricinus* density and *Borrelia burgdorferi* s.l. prevalence by combining landscape variables, forest structure, climate, and mammal community data within a single predictive model. Unlike previous studies that mostly examine these drivers in isolation, our approach reveals complex, scale-dependent interactions shaping tick-borne disease risk. Our findings support a nuanced view of the dilution effect—a hypothesis empirically supported in simpler North American systems—but still debated in the context of Europe’s more complex host communities. We show that while greater mammal species richness and a higher relative abundance of small mammals reduce *Borrelia* prevalence, increased diversity of large non-predatory mammals amplifies it.

### Landscape drivers

At the landscape scale (500 m buffer), we identified variables that reduce both tick density and *Borrelia* prevalence. A lower proportion of broadleaf forest was associated with a decrease in nymph density, likely due to the loss of favorable microhabitat conditions such as high humidity and dense leaf litter, which support tick survival and activity and reduced presence of small mammal hosts that thrive in such environments (Tack *et al*., 2012, Ruyts *et al*., 2018, Zajac *et al*., 2021). In contrast, a higher proportion of broadleaf forest led to a decrease in *Borrelia* spp. prevalence. Previously, similar results have been observed for *B. afzelii* – a small mammal-associated genospecies, potentially due to a more diverse host community in broadleaf environments, which disrupt pathogen transmission cycles by increasing the presence of non-competent reservoir hosts. This pattern is consistent with the dilution effect hypothesis (Ruyts *et al*., 2016, Hofmeester *et al*., 2017, Keesing & Ostfeld, 2021). Along the same lines, our results show that a greater forest-type diversity — reflecting landscape heterogeneity — was linked to lower *Borrelia* spp. prevalence. A similar pattern was also observed experimentally by Bourdin *et al*. (2022), though at smaller landscape scales.

Surprisingly, and in contrast to previous literature, forest cover showed a weak negative effect on overall nymph density but had no significant influence on *Borrelia* prevalence (Zajac *et al*., 2021). Similarly, forest edge density — previously associated with higher tick densities and *Borrelia* spp. prevalence (Tack *et al*., 2012, Ehrmann *et al*., 2018, Boulanger *et al*., 2024) — had no significant effects in our models when used as predictor instead of the highly collinear forest cover (data not shown).

Generalization of landscape effects should be interpreted with caution, as variable selection across spatial scales can yield differing or even opposing results (Ehrmann *et al*., 2018). Moreover, our survey plot was relatively small (300 m²), suggesting that broader landscape features may be less influential than fine-scale microhabitat conditions in shaping local tick density and *Borrelia* spp. prevalence.

### Local drivers

At the plot scale, several local forest habitat characteristics were associated with reductions in both *I. ricinus* nymph density and *Borrelia* prevalence.

We show that a higher SMI, reflecting more intensive forest use and higher conifer share, is associated with lower nymph density. This is consistent with previous findings that conifer-dominated and intensively managed forests generally support fewer ticks (Tack *et al*., 2012, Schall & Ammer, 2013, Olsthoorn *et al*., 2025). Interestingly, an earlier three-year study on tick density, conducted in the same plots within the Swabian Alb found highest tick densities in highly managed young stands (Lauterbach *et al*., 2013). However, the study also highlighted the interannual variability in the predictive strength of environmental covariates and intriguingly a comparison of the average tick density on the same 25 plots approximately 15 years later revealed no correlation whatsoever between past and present tick densities (r^2^ = 0.003, data not shown). In further contrast to previous studies, we found no direct effect of forest management intensity on *Borrelia* spp. prevalence (Olsthoorn *et al*., 2025).

The number of tree species per hectare was negatively associated with nymph density in our study, suggesting that increased tree diversity may reduce tick abundance by modifying microclimatic conditions or altering host communities—possibly through more complex food webs that decrease the availability of key tick hosts (Kahl & Gray, 2023, Vacek *et al*., 2023).

Interestingly, crown projection area, indicative of older stands with denser canopies and typically higher humidity, was negatively associated with both nymph density – albeit with a small effect – and *Borrelia* spp. prevalence. This contrasts the common assumption that closed canopies benefit ticks by enhancing humidity - a factor that, notably, did not show any effect on nymph density in our study (Tukahirwa, 1976), but it aligns with findings by Lauterbach *et al*. (2013), who observed higher nymph densities in younger stands. Furthermore, it suggests that the tick density-reducing effect of higher SMI is not primarily driven by tree felling and the resulting canopy opening. Greater shrub cover was another factor associated with lower *Borrelia* spp. prevalence, supposedly by enhancing habitats for a more diverse mammal community (Ruyts *et al*., 2016, Ehrmann *et al*., 2018).

Overall, we assume that many landscape and forest management effects discussed thus far are indirectly mediated by changes in the mammalian host community.

### Mammal host community drivers

Indeed, the mammal host community composition plays a crucial role in regulating tick density and *Borrelia* spp. prevalence, in our study.

Deer are regarded as essential for maintaining adult *Ixodes ricinus* populations, while small mammals, such as *Apodemus spp.* and *Myodes glareolus*, sustain larval stages (Hofmeester *et al*., 2017). It should be noted that most *Borrelia* genospecies exhibit host specificity. While *B. garinii* and *B. valaisiana* are found in birds, *B. afzelii* is harboured by rodents, and *B. burgdorferi* sensu stricto is associated with both host groups (Margos *et al*., 2019). This underlines the need for future research to include genospecies-level identification in relation to host community composition, particularly in complex European ecosystems with a broad range of reservoir hosts. We used camera trapping as an effective, non-invasive method for assessing mammal communities (Rovero & Zimmermann, 2016). While this method may introduce bias due to repeated sightings of the same individuals, the data also reflects host movement and activity, which are key determinants of tick encounter rates (Hofmeester *et al*., 2017).

We show that overall mammal species richness reduces nymph density and substantially lowers *Borrelia spp.* prevalence. Thus, our findings support the dilution effect hypothesis, suggesting that in the studied forest system higher mammalian species richness is associated with a decrease in the proportion of competent hosts (Keesing *et al*., 2006, Ogden & Tsao, 2009). Distinguishing between predators and prey—and among prey, between large and small species—provided more nuanced insights into the factors influencing both *I. ricinus* nymph density and *Borrelia* spp. prevalence. Large non-predatory mammals, despite including roe deer (76.88% of this group) and Leporidae (5.96%)– widely acknowledged as multipliers of tick populations – did not significantly impact nymph density in our model (Hofmeester *et al*., 2017, Boulanger *et al*., 2024, Olsthoorn *et al*., 2025). However, increased diversity among large non-predatory mammals led to higher *Borrelia* spp. prevalence, likely reflecting a lower proportion of roe deer, which act as dilution hosts (Ruyts *et al*., 2018). Predator abundance had a clear suppressive effect on nymph density, supporting findings from the Netherlands, Finland, and the USA, where intact predator communities can reduce tick densities by controlling small mammal populations and limiting their movements, reducing tick burden (Levi *et al*., 2012, Hofmeester *et al*., 2017, Terraube, 2019). As key hosts for tick larvae, small mammals, particularly *Apodemus* spp. (63.92% of small mammal species in our study) and *Myodes glareolus* (25.14%) serve as primary amplifiers of larval populations and, consequently, strong predictors of nymph density (Hofmeester *et al*., 2017). And indeed, we also found a higher diversity, though not abundance, of small mammals being associated with increased *I. ricinus* density. Intriguingly, despite mice and voles being considered competent hosts for *Borrelia spp.,* we found the relative abundance of small mammals to lower *Borrelia* prevalence (Ruyts *et al*., 2018). The absence of a direct relationship between the relative abundance of small mammals and tick density over the course of a full year aligns with findings from other European studies (Gandy *et al*., 2022, Boulanger *et al*., 2024).

### Seasonal mammal host community drivers

As described above, accounting for seasonal dynamics is essential for fully capture the effects of mammal communities on nymph density. We analysed two consecutive seasonal transitions - summer to autumn 2023 and autumn 2023 to spring 2024 - which revealed the critical role of small mammals in driving *I. ricinus* nymph density, consistent with their function as primary larval hosts (Mihalca & Sándor, 2013, Olsthoorn *et al*., 2025). Interestingly, this effect was not evident when considering the entire study period (see above) highlighting why studies over longer timescales often yield inconsistent results (Ruyts *et al*., 2018).

Our findings indicate that these relationships are more nuanced than expected. While small mammal abundance consistently predicted higher nymph density, the effect of their diversity varied across seasons. For instance, during the directly consecutive summer and autumn seasons, increased small mammal diversity was associated with lower nymph density. However, this relationship reversed in the season spanning autumn to spring. Notably, spring also showed a surprising negative association between large mammal abundance and nymph density, along with a positive correlation with predator diversity—patterns that are difficult to interpret. Possibly the unusually mild winter of 2023/2024 obscured some of the seasonal effects.

In contrast, our seasonal analysis did not reveal any clear relationship between mammal community and *Borrelia* spp. prevalence. This effect might manifest itself more long-term than the seasonal and undulating *Borrelia* spp. prevalence that lingers in the ticks (Mysterud *et al*., 2013, Hartemink *et al*., 2021).

The ongoing debate about the dilution effect hypothesis — supported by some studies (Ostfeld & Keesing, 2000, LoGiudice *et al*., 2008, Keesing *et al*., 2010, Ostfeld & Keesing, 2013, Civitello *et al*., 2015) and questioned by othersy (Chatterjee *et al*., 2012, Randolph & Dobson, 2013, Salkeld *et al*., 2013, Wood & Lafferty, 2013, Linske, 2017) — underscores the complexity of host–pathogen–habitat interactions. Given these controversial and ambiguous findings, this study, which comprehensively analyses potential influencing factors and integrates them into a single model, is crucial for moving towards a clearer conclusion.

## Conclusion

Our results, though complex given the many variables involved, reveal an intricate interplay of influencing factors, some of which remain ambiguous — particularly in the seasonal analysis. Future studies should aim for larger sample sizes to allow for methods such as Structural Equation Modeling (SEM) to disentangle the direct and indirect effects of habitat and mammal communities or even individual species on tick density and Borrelia prevalence. Nevertheless, we can identify a diluting effect of habitat and mammal host diversity: Landscapes with a diverse mix of forest types and a — not necessarily extensive — forest management that promotes a high local tree species richness, along with greater overall mammal species richness, especially with abundant predator communities, tend to support lower tick densities and reduced *Borrelia* spp. prevalence.

## Acknowledgements

We thank the local management team of the Schwäbische Alb exploratory and the managers of all three exploratories, Julia Bass, Sven Pompe, Anna K. Franke, Robert Künast, Franca Marian, Melissa Jüds, Uta Schumacher and all former managers for their work in maintaining the plot and project infrastructure; Victoria Grießmeier for giving support through the central office, Andreas Ostrowski for managing the central database, and Markus Fischer, Eduard Linsenmair, Dominik Hessenmöller, Daniel Prati, Ingo Schöning, François Buscot, Ernst-Detlef Schulze, Wolfgang W. Weisser and the late Elisabeth Kalko for their role in setting up the Biodiversity Exploratories programme. We also thank the Biodiversaty Exploratories Core Projects “Instrumentation and remote sensing”, “Botany” and “Forest structure” for providing the habitat parameters for the analysis and “Synthesis” for their initial support with statistical analysis. We thank the administration of the UNESCO Biosphere Reserve Swabian Alb as well as all land owners for the excellent collaboration. Field work permits were issued by the responsible state environmental office of Baden-Württemberg.

