## Supplemental Tables S1- S9 for "Landscape heterogeneity, forest structure and mammalian host diversity shape *Ixodes ricinus* density and *Borrelia* prevalence in German forests"

**Supplementary Material**

**S 1:** Shows the total number of flagged *Ixodes ricinus* nymphal ticks collected per site and season, as well as the number of individuals that tested positive for *Borrelia burgdorferi* sensu lato (n = positive ticks / total number of tested ticks (%; CI 95%)). Note that two ticks were excluded from testing and are therefore not included in the results.

| Spot | Spring 2023 | Summer 2023 | Autumn 2023 | Spring 2024 | Ticks total |
|---|---|---|---|---|---|
| AEW 1 | 0/0 | 0/5 (0.0%, 0.0-52.2%) | 0/4 (0.0%, 0.0-60.2%) | 0/1 (0.0%, 0.0-97.5%) | 0/10 (0.0%, 0.0-30.9%) |
| AEW 2 | 2/11 (18.2%, 2.3-51.8%) | 2/14 (14.3%, 1.8-42.8%) | 0/3 (0.0%, 0.0-70.8%) | 2/13 (15.4%, 1.9-45.5%) | 6/42 (14.3%, 5.4-28.5%) |
| AEW 3 | 0/2 (0.0%, 0.0-84.2%) | 0/1 (0.0%, 0.0-97.5%) | 1/9 (11.1%, 0.3-48.2%) | 0/4 (0.0%, 0.0-60.2%) | 1/16 (6.2%, 0.2-30.2%) |
| AEW 4 | 0/8 (0.0%, 0.0-36.9%) | 2/17 (11.8%, 1.5-36.4%) | 0/10 (0.0%, 0.0-30.9%) | 0/6 (0.0%, 0.0-45.9%) | 2/42 (4.8%, 0.6-16.2%) |
| AEW 5 | 0/18 (0.0%, 0.0-18.5%) | 0/8 (0.0%, 0.0-36.9%) | 0/27 (0.0%, 0.0-12.8%) | 1/39 (2.6%, 0.1-13.5%) | 1/92 (1.1%, 0.0-5.9%) |
| AEW 6 | 1/42 (2.4%, 0.1-12.6%) | 3/36 (8.3%, 1.8-22.5%) | 3/11 (27.3%, 6.0-61.0%) | 0/21 (0.0%, 0.0-16.1%) | 7/110 (6.4%, 2.6-12.7%) |
| AEW 7 | 1/26 (3.8%, 0.1-19.6%) | 1/25 (4.0%, 0.1-20.3%) | 4/13 (30.8%, 9.1-61.4%) | 1/15 (6.7%, 0.2-31.9%) | 7/79 (8.9%, 3.6-17.4%) |
| AEW 8 | 3/13 (23.1%, 5.0-53.8%) | 1/19 (5.3%, 0.1-26.0%) | 0/2 (0.0%, 0.0-84.2%) | 2/140 (1.4%, 0.2-5.1%) | 6/174 (3.4%, 1.3-7.3%) |
| AEW 9 | 3/28 (10.7%, 2.3-28.2%) | 4/26 (15.4%, 4.4-34.9%) | 1/4 (25.0%, 0.6-80.6%) | 0/5 (0.0%, 0.0-52.2%) | 8/63 (12.7%, 5.7-23.5%) |
| AEW 11 | 0/2 (0.0%, 0.0-84.2%) | 0/3 (0.0%, 0.0-70.8%) | 0/3 (0.0%, 0.0-70.8%) | 0/2 (0.0%, 0.0-84.2%) | 0/10 (0.0%, 0.0-30.9%) |
| AEW 12 | 1/6 (16.7%, 0.4-64.1%) | 1/5 (20.0%, 0.5-71.6%) | 0/3 (0.0%, 0.0-70.8%) | 0/7 (0.0%, 0.0-41.0%) | 2/21 (9.5%, 1.2-30.4%) |
| AEW 13 | 0/2 (0.0%, 0.0-84.2%) | 0/13 (0.0%, 0.0-24.7%) | 1/6 (16.7%, 0.4-64.1%) | 0/10 (0.0%, 0.0-30.9%) | 1/31 (3.2%, 0.1-16.7%) |
| AEW 14 | 0/7 (0.0%, 0.0-41.0%) | 0/6 (0.0%, 0.0-45.9%) | 1/2 (50.0%, 1.3-98.7%) | 0/1 (0.0%, 0.0-97.5%) | 1/16 (6.2%, 0.2-30.2%) |
| AEW 17 | 1/5 (20.0%, 0.5-71.6%) | 0/10 (0.0%, 0.0-30.9%) | 0/2 (0.0%, 0.0-84.2%) | 0/7 (0.0%, 0.0-41.0%) | 1/24 (4.2%, 0.1-21.1%) |

| | | | | | |
|---|---|---|---|---|---|
| AEW 18 | 0/23 (0.0%, 0.0-14.8%) | 1/14 (7.1%, 0.2-33.9%) | 0/10 (0.0%, 0.0-30.9%) | 9/56 (16.1%, 7.6-28.3%) | 10/103 (9.7%, 4.8-17.1%) |
| AEW 20 | 0/10 (0.0%, 0.0-30.9%) | 0/9 (0.0%, 0.0-33.6%) | 0/10 (0.0%, 0.0-30.9%) | 1/19 (5.3%, 0.1-26.0%) | 1/48 (2.1%, 0.1-11.1%) |
| AEW 23 | 0/12 (0.0%, 0.0-26.5%) | 0/29 (0.0%, 0.0-11.9%) | 4/47 (8.5%, 2.4-20.4%) | 6/22 (27.3%, 10.7-50.2%) | 10/110 (9.1%, 4.5-16.1%) |
| AEW 31 | 0/1 (0.0%, 0.0-97.5%) | 0/2 (0.0%, 0.0-84.2%) | 1/6 (16.7%, 0.4-64.1%) | 0/12 (0.0%, 0.0-26.5%) | 1/21 (4.8%, 0.1-23.8%) |
| AEW 38 | 3/25 (12.0%, 2.5-31.2%) | 1/35 (2.9%, 0.1-14.9%) | 0/7 (0.0%, 0.0-41.0%) | 2/12 (16.7%, 2.1-48.4%) | 6/79 (7.6%, 2.8-15.8%) |
| AEW 39 | 0/5 (0.0%, 0.0-52.2%) | 2/9 (22.2%, 2.8-60.0%) | 1/3 (33.3%, 0.8-90.6%) | 2/9 (22.2%, 2.8-60.0%) | 5/26 (19.2%, 6.6-39.4%) |
| AEW 40 | 0/0 | 2/20 (10.0%, 1.2-31.7%) | 0/6 (0.0%, 0.0-45.9%) | 0/1 (0.0%, 0.0-97.5%) | 2/27 (7.4%, 0.9-24.3%) |
| AEW 42 | 0/56 (0.0%, 0.0-6.4%) | 0/27 (0.0%, 0.0-12.8%) | 2/28 (7.1%, 0.9-23.5%) | 0/15 (0.0%, 0.0-21.8%) | 2/126 (1.6%, 0.2-5.6%) |
| AEW 43 | 0/0 | 0/11 (0.0%, 0.0-28.5%) | 0/2 (0.0%, 0.0-84.2%) | 0/9 (0.0%, 0.0-33.6%) | 0/22 (0.0%, 0.0-15.4%) |
| AEW 49 | 0/3 (0.0%, 0.0-70.8%) | 2/11 (18.2%, 2.3-51.8%) | 3/26 (11.5%, 2.5-30.1%) | 2/9 (22.2%, 2.8-60.0%) | 7/49 (14.3%, 5.9-27.2%) |
| AEW 50 | 1/33 (3.0%, 0.1-15.8%) | 0/30 (0.0%, 0.0-11.6%) | 2/10 (20.0%, 2.5-55.6%) | 2/25 (8.0%, 1.0-26.0%) | 5/98 (5.1%, 1.7-11.5%) |
| Total | 16/338 (4.7%, 2.7-7.6%) | 22/385 (5.7%, 3.6-8.5%) | 24/254 (9.4%, 6.2-13.7%) | 30/462 (6.5%, 4.4-9.1%) | 92/1437 (6.4%, 5.2-7.8%) |

**S 2**: Conditional averaged generalized linear regression results using a Gamma distribution with a logarithmic link function for effects of silvicultural management index (SMI), relative abundance index (RAI), Shannon' diversity index (H) and species richness (S) of mammals, local forest structure (CPA = crown projection area) and landscape variables on nymph density. Significant variables are shown in bold.

| **Predictor** | **Estimate** | **Std. Error** | **p-value** |
|---|---|---|---|
| (Intercept) | 3.561 | 1.046 | **< 0.001** |
| Broad leaf share | 2.600 | 0.569 | **< 0.001** |
| Forest cover | -1.415 | 0.604 | **0.028** |
| RAI Predators | -0.008 | 0.002 | **< 0.001** |

| | | | |
|---|---|---|---|
| SMI | -3.041 | 0.623 | **< 0.001** |
| Tree species/ha | -0.098 | 0.031 | **0.003** |
| H large mammals | 0.751 | 0.407 | 0.078 |
| H small mammals | 0.950 | 0.368 | **0.017** |
| S total mammals | -0.207 | 0.060 | **0.001** |
| CPA | -0.0001 | < 0.001 | **0.017** |

**S 3**: AICc table for the candidate models describing nymph density, displaying the models with delta AICc<2 (Full model: nymph density ~ broad leaf share + H forest + forest cover + SMI + tree species richness + crown projection area + mean tree dbh + dead wood volume + shrub cover + relative air humidity + RAI predators + H predators + RAI small mammals + H small mammals + RAI large mammals + H large mammals + S total mammals). SMI = silvicultural management index, RAI = relative abundance index, H = Shannon's diversity index, S = species richness.

| **Candidate models - nymph density** | **AICc** | **Delta AICc** | **Weight** |
|---|---|---|---|
| Intercept + broad leaf share + forest cover + SMI + tree species richness + RAI predators | 105.1 | 0.00 | 0.046 |
| Intercept + broad leaf share + forest cover + tree species richness + SMI + RAI predators + H large mammals | 106.6 | 1.49 | 0.022 |
| Intercept + broad leaf share + SMI + tree species richness + crown projection area + RAI predators + H small mammals + H large mammals + S total mammals | 106.8 | 1.66 | 0.020 |
| Intercept + broad leaf share + SMI + tree species richness + RAI predators + S total mammals | 107.0 | 1.88 | 0.018 |

**S 4**: Conditional averaged linear regression results using a normal distribution for effects of silvicultural management index (SMI), Shannon's diversity index (H), relative abundance index

(RAI) and species richness (S) of mammals, local forest structure (CPA = crown projection area, dbh = diameter at breast height) and landscape variables on *Borrelia* spp. prevalence. Significant variables are shown in bold.

| **Predictor** | **Estimate** | **Std. Error** | **p-value** |
|---|---|---|---|
| (Intercept) | 118.000 | 28.080 | **<0.001** |
| Broad leaf share | -17.590 | 5.279 | **0.002** |
| H Forest | -10.250 | 3.825 | **0.012** |
| Rel. humidity | -0.937 | 0.303 | **0.004** |
| Shrub cover | -0.089 | 0.039 | **0.032** |
| CPA | -0.002 | 0.0007 | **0.007** |
| H large mammals | 8.663 | 4.155 | **0.045** |
| RAI small mammals | -0.001 | 0.0004 | **0.014** |
| S total mammals | -1.245 | 0.483 | **0.017** |
| Mean tree dbh | 0.147 | 0.088 | 0.122 |
| Forest cover | 9.218 | 5.248 | 0.104 |

**S 5**: AICc table for the candidate models describing *Borrelia spp.* prevalence, displaying the models with delta AICc<2 (Full model: *Borrelia* spp. prevalence ~ broad leaf share + H forest + forest cover + SMI + tree species richness + crown projection area + mean tree dbh + dead wood volume + shrub cover + relative air humidity + RAI predators + H predators + RAI small mammals + H small mammals + RAI large mammals + H large mammals + S total mammals). H = Shannon's diversity index, RAI = relative abundance index, S = species richness, dbh = diameter at breast height.

| **Candidate models – *Borrelia* spp. prevalence in nymphs** | **AICc** | **Delta AICc** | **Weight** |
|---|---|---|---|
| Intercept + broad leaf share + H forest + shrub + crown projection area + relative air humidity | 146.9 | 0.00 | 0.024 |

| | | | |
|---|---|---|---|
| Intercept + broad leaf share + H forest + crown projection area + relative air humidity + RAI small mammals + H large mammals + S total mammals | 147.6 | 0.61 | 0.017 |
| Intercept + broad leaf share + H forest + mean tree dbh + shrub cover + crown projection area + relative air humidity | 147.8 | 0.86 | 0.015 |
| Intercept + broad leaf share + H forest + shrub cover + crown projection area + relative air humidity + H large mammals | 148.7 | 1.78 | 0.010 |
| Intercept + broad leaf share + H forest + forest cover + crown projection area + relative air humidity + RAI small mammals + H large mammals + S total mammals | 148.9 | 1.92 | 0.009 |
| Intercept + broad leaf share + H forest + crown projection area + relative air humidity | 148.9 | 1.98 | 0.009 |

**S 6**: Best fitted generalized linear regression model using a Gamma distribution with a logarithmic link function for effects of relative abundance index (RAI) and Shannon diversity (H) of mammals in the preceding seasons on nymph density in spring and autumn. Significant variables are shown in bold.

| **Predictor** | **Estimate** | **Std. Error** | **p-value** |
|---|---|---|---|
| **Autumn '23** | | | |
| (Intercept) | 0.857 | 0.285 | **0.006** |
| RAI small mammals | 0.008 | 0.002 | **<0.001** |
| H small mammals | -1.236 | 0.460 | **0.014** |
| **Spring '24** | | | |
| (Intercept) | 0.298 | 0.628 | 0.640 |
| RAI small mammals | 0.0004 | <0.001 | **0.012** |
| RAI large mammals | -0.272 | 0.010 | **0.010** |
| H small mammals | 1.873 | 0.668 | **0.020** |
| H predators | 1.731 | 0.681 | **0.011** |

**S 7**: AICc table for the candidate models describing nymph density in autumn 2023 and spring 2024, displaying the models with delta AICc<2 (Full models: nymph density ~ RAI predators + H predators + RAI small mammals + H small mammals + RAI large mammals + H large mammals). H= Shannon's diversity index, RAI = relative abundance index.

| **Candidate models – nymph density autumn '23** | **AICc** | **Delta AICc** | **Weight** |
|---|---|---|---|
| Intercept + RAI small mammals + H small mammals | 98.4 | 0.00 | 0.357 |
| **Candidate models – nymph density spring '24** | | | |
| Intercept + H Predators + RAI small mammals + H small mammals + RAI large mammals | 139.5 | 0.00 | 0.196 |

**S 8**: Conditional averaged generalized linear regression results using a tweedie distribution for effects of relative abundance index (RAI) and Shannon's diversity index (H) of mammals in the preceding seasons on *Borrelia* spp. prevalence. Significant variables are shown in bold.

| Predictor | Estimate | Std. Error | p-value |
|---|---|---|---|
| **Autumn '23** | | | |
| (Intercept) | 2.426 | 0.338 | **<0.001** |
| RAI Predators | -0.033 | 0.033 | 0.348 |
| **Spring '24** | | | |
| (Intercept) | 1.010 | 0.767 | 0.205 |
| H large mammals | 1.903 | 1.135 | 0.114 |
| RAI large mammals | 0.016 | 0.011 | 0.185 |
| H predators | 1.161 | 0.922 | 0.235 |

**S 9**: AICc table for the candidate models describing *Borrelia* spp. prevalence in autumn 2023 and spring 2024, displaying the models with delta AICc<2 (Full models: *Borrelia* spp. prevalence ~ RAI predators + H predators + RAI small mammals + H small mammals + RAI large mammals + H large mammals). H= Shannon’s diversity index, RAI = relative abundance index.

| **Candidate models – *Borrelia* spp. prevalence autumn ‘23** | **AICc** | **Delta AICc** | **Weight** |
|---|---|---|---|
| Intercept | 132.1 | 0.00 | 0.234 |
| Intercept + RAI predators | 133.8 | 1.77 | 0.097 |
| **Candidate models – *Borrelia* spp. prevalence spring ‘24** | | | |
| Intercept + H large mammals | 117.9 | 0.00 | 0.122 |
| Intercept | 118.3 | 0.36 | 0.102 |
| Intercept + RAI large mammals | 118.7 | 1.80 | 0.082 |
| Intercept + H predators | 119.6 | 1.67 | 0.053 |
| Intercept + RAI large mamals + H large mammals | 119.8 | 1.88 | 0.048 |
